# Responding kinetic of B-cell receptor repertoire to the Toll-like receptor 7/8 stimulation in non-human primates

**DOI:** 10.1101/2021.03.15.435360

**Authors:** Shiyu Wang, Judith Mandl, Mark Feinberg, Michael Citron, Nitin K. Saksena, Lihua Luo, Zida Li, Yonggang Zhu, Tao Li, Chao Nie, Xiuqing Zhang, Ya Liu, Xiao Liu, I-Ming Wang, Wei Zhang

## Abstract

TLR7 and 8 regulate B cell immunity, but the precise details of the mechanism are still unclear. Here, we studied the kinetics of both heavy and light chains (IgKL) of B-cell receptor (BCR) repertoire responding to the TLR7/8 stimulation in two geniuses of non-human primates (NHPs), African green monkeys (AGMs) and rhesus macaques (RMs). We evaluated the activation of lymphocytes by flow cytometry, and studied characteristics of BCR repertoire in terms of gene usage, repertoire diversity, and the number of lineages. Although AGMs had a weaker activation than RMs, and a different responding kinetic, both AGMs and RMs presented an increased IgKL repertoire diversity and lineages expansion. It suggested that the responding time rather than initiation of TLR7/8-induced IgKL repertoire response related to B cell activation. Expanded IgKL lineages with frequency from 0.001% to 1% had an elevated mutation rate and expanded IgH lineages used more IgA/G/E, suggesting that the TLR7/8 stimulation expanded low-frequent but high-mutated lineages. Besides, most of expanded IgKL lineages were λ isotype. In conclusion, TLR7/8 selectively expands IgKL lineages with a high mutation rate, low frequency, and λ isotype. The selective effect of TLR7/8 on BCR repertoire allows TLR7/8 agonists to be adjuvant for selectively accelerating antibody maturation.

## Introduction

Efforts to induce neutralizing antibodies by vaccination are effective methods to prevent virus infections. However, vaccinations often meet disappointing results for unclear reasons, such as HIV vaccinations. Early during HIV infection, antibodies to gp120 are produced but failed to block virus. The high mutation rate of the envelope glycoprotein is a contributing factor. Evenly, some individuals generate broadly neutralizing antibodies, but these arise too late to be of clinical beneficial^1^. Vaccines for other viruses, such as influenza, also meet a similar problem. Thereby, accelerating the generation of antibodies with a high quality is a key priority^2^.

Over the past decade, it has become clear that innate immune system contributes to the activation and regulation of adaptive immune response, although the precise details are still unclear. Innate immune system senses the presence of viral particles by three classes of sensors, NOD like receptors, RIG-I like receptors, and Toll like receptors (TLRs). TLRs affected the antibody maturation at stages of B cell activation, germinal center formation, antibody somatic hypermutation (SHM), and antibody class switching^3^. TLR7 and the closely related TLR8 are pattern recognition receptors for purine-rich single-stranded ribonucleic acid (ssRNA)^4^, which is common in retroviruses, such as HIV. Stimulations via TLR7 and 8 can promote immune response trigged by vaccination. A combination of TLR7 agonist and inactivated influenza virus accelerated antigen-specific antibody maturation. The underlying mechanism may basically fall in two classes. Firstly, Simulations via TLR7/8 in the plasmacytoid dendritic cells (pDCs)^5,6^ lead production of proinflammatory cytokines, such as type one interferons, to activate other immune cells, including T and B lymphocytes^7^. Secondly, TLR7 constitutively expressed in B cells regulates a key checkpoint controlling development of germinal center B cells and antibody maturation. In mice, deletion of Myd88 in B cells rather than DCs disrupts immune control of Friend virus. Recently, TLR7 pathway is shown to combinate with B-cell receptor (BCR) pathway to shape the antibody maturation processes in B cells. The TLR7 activation facilitate the pathway of BCR special to self-antigens in IgM memory B cells, and B1-like cells^8^. However, these conclusions are unstable among situations. Part of reasons should be the combinations of vaccine and TLR7/8 agonist, since components of vaccines can play adjuvant roles, and the effect of TLR7/8 will be masked by effects of vaccines.

During the past decade, BCR sequencing (BCR-seq) promote our knowledge of the humoral immunity. Studies employing this technique unveil antibody evolution in human and animal models after vaccination and infections. Scott Boyd and his colleges described the kinetic of class-switching and somatic hyper mutation (SHM) of antibody responses to Ebola virus infection^9^. An *in vitro* study using BCR-seq showed that the TLR7/8 stimulation alone biased to expand IgM BCR^10^, and therefore may skewed the BCR repertoire *in vivo*. Therefore, studies on BCR repertoire responses to the TLR7/8 stimulation will give a new view to understand how TLR7/8 affect B cell immunity *in vivo*.

Non-human primates (NHPs) are suitable models for simulating immune responses of human beings to vaccines and treatments^11^. Rhesus macaques (RM) and African green monkeys (AGM) were used as a comparison to study pathology of simian immunodeficiency virus (SIV) and vaccination mechanism. These studies showed that immune response of AGMs to TLRs stimulation was weaker than that of RMs, such as a lower type one interferon production in AGMs^12–14^. Therefore, we employed RMs and AGMs as a comparison to unveil how TLR7/8 affected BCR repertoire when TLR7/8 signaling pathways were different.

Immune response during the first week is important for a vaccine to provide an effective protection^15,16^. We collected samples within one-week post-stimulation to uncover the TLR7/8 stimulation’s influence during this period. Since TLR9 is also expressed in B cells and affects B cell activation^7,17,18^, we stimulated these NHPs with a TLR9 agonist two months later following the same scheme as a contrast.

## Methods and materials

### Agonists

IDR-053 is a synthetic TLR7/TLR8 (TLR7/8) agonist with sequence 5’-X5UGCUGCUUGUG-X-GUGUUCGUCGUX5-5’, where X is a glycerol linker and X5 represents 1, 5 pentanediol, the precise method to synthesize this molecule is recorded in patent EP2357231A2. IMO-2125 is a synthetic TLR9 agonist. These two compounds were synthesized as previously described^19^.

### Animals, injections and sample collection

Six disease-free rhesus macaques (*Macaca mulatta*) and six African green monkeys (*Chlorocebus aethiops*) were employed in this study (The detailed records per NHP were listed in **Supplementary Table 1**). All NHPs were housed at New Iberia Research Center in a system by adhering to standards of the Guide for the Care and Use of Laboratory Animals, the Public Health Service Policy on Humane Care and Use of Laboratory Animal, AVMA Guidelines for the Euthanasia of Laboratory Animals, the USDA Animal Welfare Act and Regulations.

Peripheral blood was collected pre-stimulation as a baseline. After dosing 25mg IDR-053 or 2.5 mg IMO-2125, blood was collected at 8hrs, 24hrs, 48hrs, 72hrs, and 1-week post-stimulation. Blood samples (~ 2.6 ml) collected at pre-and-post-vaccination at 24, 72hrs and 1-week were analyzed by fluorescence-activated cell sorting (FACS). Peripheral blood mononuclear cells (PBMCs) were collected from blood samples (~ 2ml) at all time-points with Ficoll-Paque (GE Healthcare) centrifuge method.

### Flow cytometry

For cell surface staining, fresh PBMCs were resuspended in 100 μl of flow cytometry staining buffer (3% BSA in PBS) containing primary antibody ligated with biotin against CD3 (Supplementary Table2), incubated for 40 min at 4°C and were washed with FACS staining buffer. Then cells were resuspended in 100ul of flow cytometry staining buffer containing antibodies against biotin, CD20 and HLA-DR (Supplementary Table2) and were incubated for 40 min at 4°C. For intracellular staining, cells were washed with flow cytometry staining buffer, permeabilized with cytofix/cytoperm Kit (BD Biosciences, Cat. no: 554714), and then stained with anti-Ki-67 antibody. The stained cells were analyzed by a BD FACS Aria II cell sorter (BD Biosciences). B cells were gated by CD3^-^/CD20^low/+^.

### Library preparation and sequencing

BCR repertoire library was prepared with RNA (600 micrograms) extracted from PBMCs using Trizol (Invitrogen,15596-026) and 5’ Rapid amplification of complementary DNA (cDNA) ends (5’RACE, Invitrogen, 18374-058) method as described previously^20,21^. In brief, the first-strand cDNA was synthesized using SuperScript II reverse transcriptase (Invitrogen), an abridged anchor primer (AAP), as an upstream primer, and a pool of downstream primers for IGH C-gene labeled with biotin, including the four published in previous study^20^., along with the eleven newly designed for the purpose of this study (**supplementary table 2**). Following that, the BCRs were amplified by polymerase chain reaction (PCR) with a forward primer annealing to the AAP region and a pool of reverse primers annealing to the C region; PCR cycling conditions were 94°C for 2 min, 30 cycles of 94°C for the 30s, 60°C for 30s and 72°C for 45 s, and 72°C for 7 min, followed by the final step at 4°C as a holding temperature. PCR products were purified with Ampure XP beads (Agencourt) and fragmented into 150-200bp using Covaris E220; fragments containing a biotinylated C region were captured by M270 beads which were crosslinked to streptavidin (Invitrogen), with a dissociation, end repairing and linkage, targeting fragments were linked to adapters and then amplified by PCR. Following qualification by Agilent 2100 Bioanalyzer system (DNA 12,000 kit) and Applied Biosystems qPCR Bioanalyzer system, samples were pooled evenly, libraries were prepared using the paired-end 150 bp kit as per manufacturer’s instruction and the library was sequenced on Hiseq2000 (Illumina) system.

### Data availability

The data supporting the findings presented in this study have already been deposited in the CNSA (https://db.cngb.org/cnsa/)^22^ of CNGBdb^23^ under the accession code CNP0000866.

### BCR sequencing data analysis

Raw data were aligned and annotated using */Monitor*^24^, as previously described. In brief, 1) sequences were quality controlled, and the qualified, paired sequences were merged; 2) BLAST was used to align sequence to IMGT (http://www.imgt.org) references, and then a re-alignment process was employed to improve the accuracy of alignment; 3) to reduce PCR and sequencing errors, nucleotide sequences for each sample were sequenced in at least 5 replicates and translated into amino acid sequences. Thus, each sequence used in BCR sequence data analysis represents an average of minimum of 5 sequences.

### Individual, sample, Clone definition

We termed a NHP as an individual. Since six RMs and six AGMs were employed, there were 12 individuals in our study. For a given individual, one blood sample was termed a sample. For a sample, a clone was defined as a unique CDR3 sequence. Abundance was termed as the times of a clone sequenced in a sample.

### Lineages definition

To increase affinity for a given antigen, BCR clones experience somatic hypermutation (SHM). These sequences with the same V- and J-gene and a single point mutation in the CDR3 region may derivate from a common ancestral cell^25^. We clustered sequences from all samples of each individual, and termed sequences using the same V-gene, J-gene and encoding CDR3 amino acid sequence with <1 mismatch as a lineage.

### Diversity and clonality evaluation

We sub-sampled 500,000 light chain sequence and 400,000 heavy chain sequence, clustered lineages, and then calculated the diversity and clonality index of unique CDR3s and lineages^26^.

Shannon entropy formula:

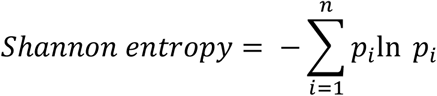

Where *p_i_* is the frequency of CDR3/lineage *i* in a sample with a size of unique CDR3/lineage *n.*

Top100 formula:

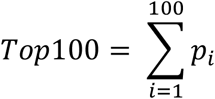

where all lineages were ranked by frequency, and p_i_ is the frequency of *i*th top lineage in one sample.

### Morisita-Horn similarity index

The frequency distribution of CDR3s from two different individuals was used to calculate the inter-individual Morisita-Horn similarity index^27^ for expanded and randomly sampled lineages.

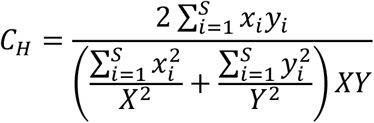

Where *X_i_* is the number of times of *i*th CDR3 in one sample with size X; *y_i_* is the number of times of *i*th CDR3 in the other sample with size Y

### Mutation rate

To account the mutation rate of a given lineage, we firstly sum mismatches in V and J regions of all amino acids sequences of the lineage by aligning these sequences to references. After that, we divided the summed number of mismatches by the number of all amino acids of both V and J regions, and then termed the ratio as the mutation rate of the lineage:

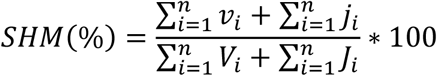

where *v_i_* is the number of mismatches in the V region with amino acid size *V_i_* in sequence *i*; j_i_ is the number of mismatches in J region with amino acids size *J_i_* in sequence *i*.

### Definition of lineage types

At a given timepoint post-stimulation, we classed lineages into five subgroups depending on their frequency, unique clone number, and mutation a lineage (three features) at this time-point and 0hr. The five subgroups were stable-lineage, increased-lineage, decreased-lineage, new-lineage, and disappeared-lineage. These definitions were performed for three features separately. For example, a lineage might be a stable-lineage by the threshold on frequency, but was classed as an increased-lineage by the threshold on mutation rate. The thresholds on each feature are shown in following. For samples of an individual, a lineage presented at 0hr but disappeared at a given time point, we termed this lineage as a disappeared-lineage at this time-point; in opposite, a lineage presented at a given time point but not presented at 0hr, we termed it as a new-lineage. For a lineage presented both at 0hr and a given time-point: in the analysis of frequency, if its frequency raised, and the increased fold change over 1, we termed it as an increased-lineage; if its frequency reduced, and the decreased fold change over 1, we termed it as a decreased-lineage; if the changed fold change of its frequency was less than 1, we termed it as a stable-lineage. For lineages presented both at 0hr and a given time-point, similar thresholds were also performed on unique clone number and mutation rate. According to previous studies, without combining with signaling via BCR, TLRs signaling only affected the B cells proliferation. Therefore, new-lineages were just a subgroup of increased-lineages defined on frequency, and disappeared-lineage were a subgroup of decreased-lineages.

### Expanded lineages identification

The enriched lineages in the peripheral blood were defined as lineages with a mutation rate <14, detected at both 48 and 72hrs post-stimulation, meeting at least one of the following requirements for the light chain lineages, 1) increased lineages with frequency > 0.001% and a fold change > 3 at 72 hours; 2) increased lineages with unique CDR3 number > 5 and a fold change > 2 at 72 hours; 3) increased lineages with mutation rate > 6 and a fold change > 1.3 at 72 hours; for heavy chain lineages: 1) increased lineages with frequency > 0.001% and a fold change > 3 at 72 hours; 2) increased lineages with mutation rate > 5 and a fold change > 1.5 at 72 hours.

### Control lineage definition

To provide quality control and assurance for evaluating features of enriched lineages, we randomly sampled lineages other than enriched ones with a matched frequency distribution at 72hrs post-stimulation with TLR7/8 agonist. Similarity and mutation rates were estimated and compared for enriched and randomly sampled lineages.

### Statistics

In this study, we used paired Wilcox-ranked tests to examine the difference between two groups (where p< 0.05 was indicated as significance) and correct p-values in multiple tests with the false discovery rate (FDR) method; correlation between two groups was estimated by Spearman’s correlation. All statistics were performed with in-house programs coded in Perl (v5.26.2) and R (3.6.0) with packages ggplot2 and dplyr.

### Ethics statement

All experiments involving laboratory animals were approved by the Merck Institutional Animal Care and Use Committee (IACUC) of Merck & Co., Inc., Kenilworth, NJ USA and Institutional Review Board (IRB) of BGI-Shenzhen (No. 13040), and were conducted in accordance with all the animal care and use laws, regulations and guidelines.

## Results

### The stimulation via TLR7/8 activated T and B lymphocytes

We stimulated six RMs and six AGMs with 25 mg TLR7/8 agonist per individual, and then collected peripheral blood from all monkeys at 6-time points, 0 hours (0hrs, pre-stimulation), 8hrs, 24hrs, 48hrs, 72hrs, and 1-week post-stimulation (**Fig. 1A**). To verify that agonist activated lymphocytes, we examined the activation markers (HLA-DR) of B cells, and activation markers (CD69) of T cells at 0hrs, 24hrs, 72hrs, and 1-week post-stimulation by flow cytometry. For B lymphocytes in both RMs and AGMs, the mean fluorescence intensity (MFI) of HLA-DR increased after stimulating, and then reached the peak at 24hrs, at last reduced from 72hrs to 1 week. Similarly, CD4^+^ T cells and CD8^+^ T cells expressed CD69 after stimulation, peaked at 24hrs, and then reduced after 72hrs (**Fig. 1B**). The increased expression of HLA-DR and CD69 suggested that TLR7/8 stimulation did activate adaptive immune cells. We also examined the proliferative marker (Ki67) of T and B lymphocytes. The percentage of Ki67^+^ in B cells, CD4^+^ T cells, and CD8^+^ T cells in RMs raised at 24hrs, and then reduced after 72hrs. However, the TLR7/8 stimulation did not lead a great change in the percentage of Ki67^+^ in CD4^+^ T, CD8^+^ T and B lymphocytes in AGM during one week (**Fig. 1C**), suggesting that the TLR7/8 stimulation promoted lymphocytes proliferation only in RMs.

**Fig.1.**
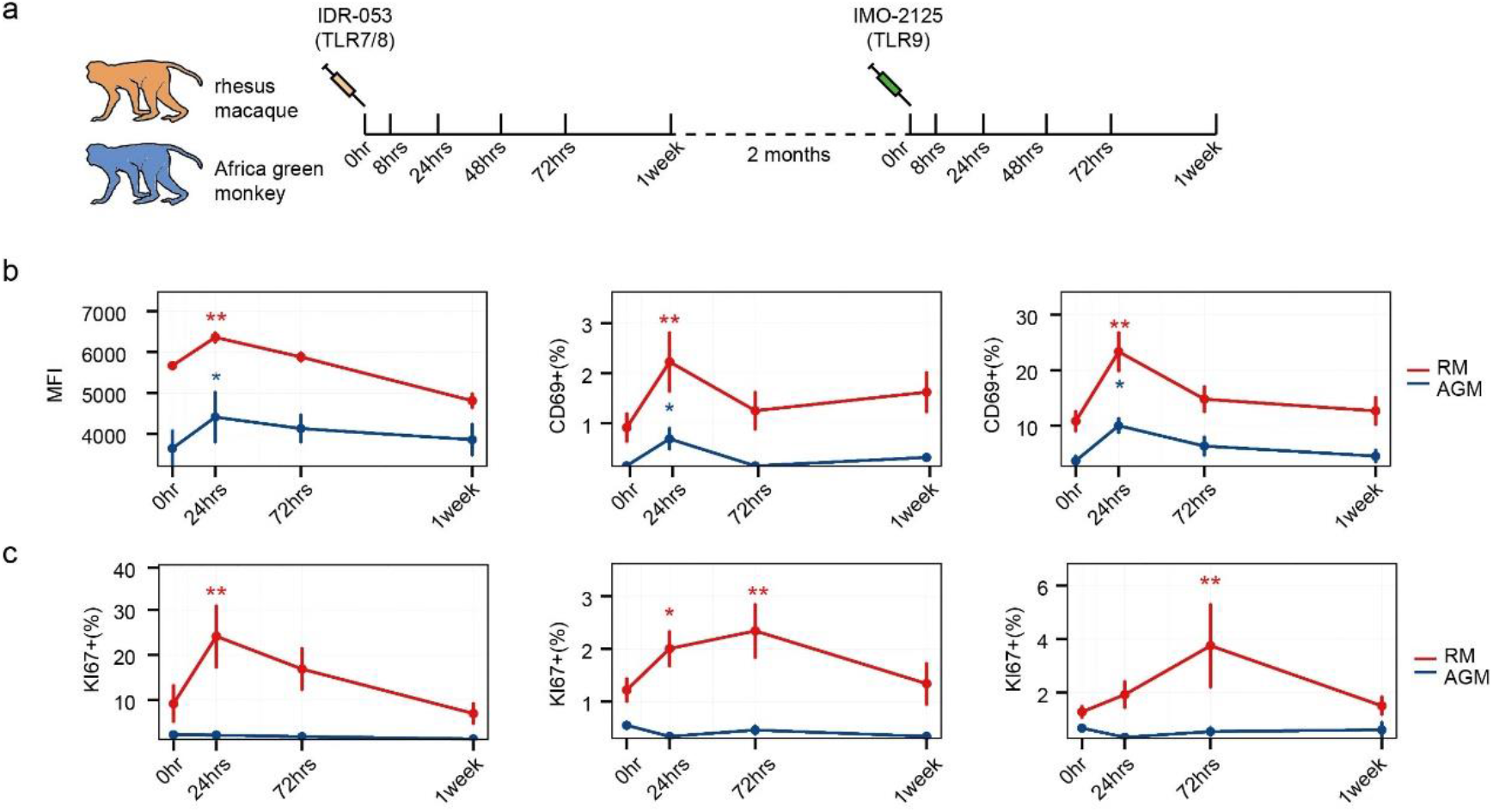
Flowchart, and activation and proliferation of T and B lymphocytes. (**a**) The TLR agonists injection and the timeline of blood collection. (**b**) the activation of B cells (MFI of HLA-DR, left), and the activation of CD4^+^ T cells (CD69^+^ percentage, median) and CD8^+^ T cells (CD69^+^ percentage, right) detected by flow cytometry. (**c**) the proliferation (Ki67 %) of B cells (left), CD4^+^ T cells (median) and CD8^+^ T cells (right) detected by flow cytometry. Paired Wilcox-ranked test was used to examine the difference between samples at 0hr and a given time-point (*p<0.05, **p<0.05).

We used 5’RACE method to amplify BCR sequences from heavy chain (IgH), and λ and κ light chains (IgKL), respectively. Because most AGM germline alleles are still unclear, it is hard to design available primers to enrich AGM heavy chain sequences. This limitation leaded a low alignment rate of AGM IgH (< 30%). Therefore, we did not study IgH repertoire of AGMs in following.

### The TLR7/8 stimulation skewed gene usage and increased BCR repertoire diversity in both RMs and AGMs

To unveil the influence of TLR7/8 stimulation on BCR repertoire, we examined the gene usage and diversity of BCR repertoire. For RMs, Gini-index of V and J gene of IgKL slightly raised at 8hrs, and then reduced gradually from 8hrs to 72hrs. After 72hours, Gini-index of V and J genes did not change significantly. For AGMs, Gini-index gradually decreased from 0 to 48hrs, and then raised after 48hrs (**Fig. 2A**). Furthermore, most of the greatly expanded genes at 72hrs were λ isotypes (**Fig. 2B** and **Fig. S1**). Notably, for AGMs, the κ to λ ratio (κ/λ) decreased from 0 to 48hrs, and then raised after 48hrs; for RMs, κ/λ declined from 0 to 72hrs, and then keep constant from 72hrs to 1 week (**Fig. 2C**). These results confirmed that λ chain genes usage indeed expanded by the TLR7/8 stimulation.

**Fig.2.**
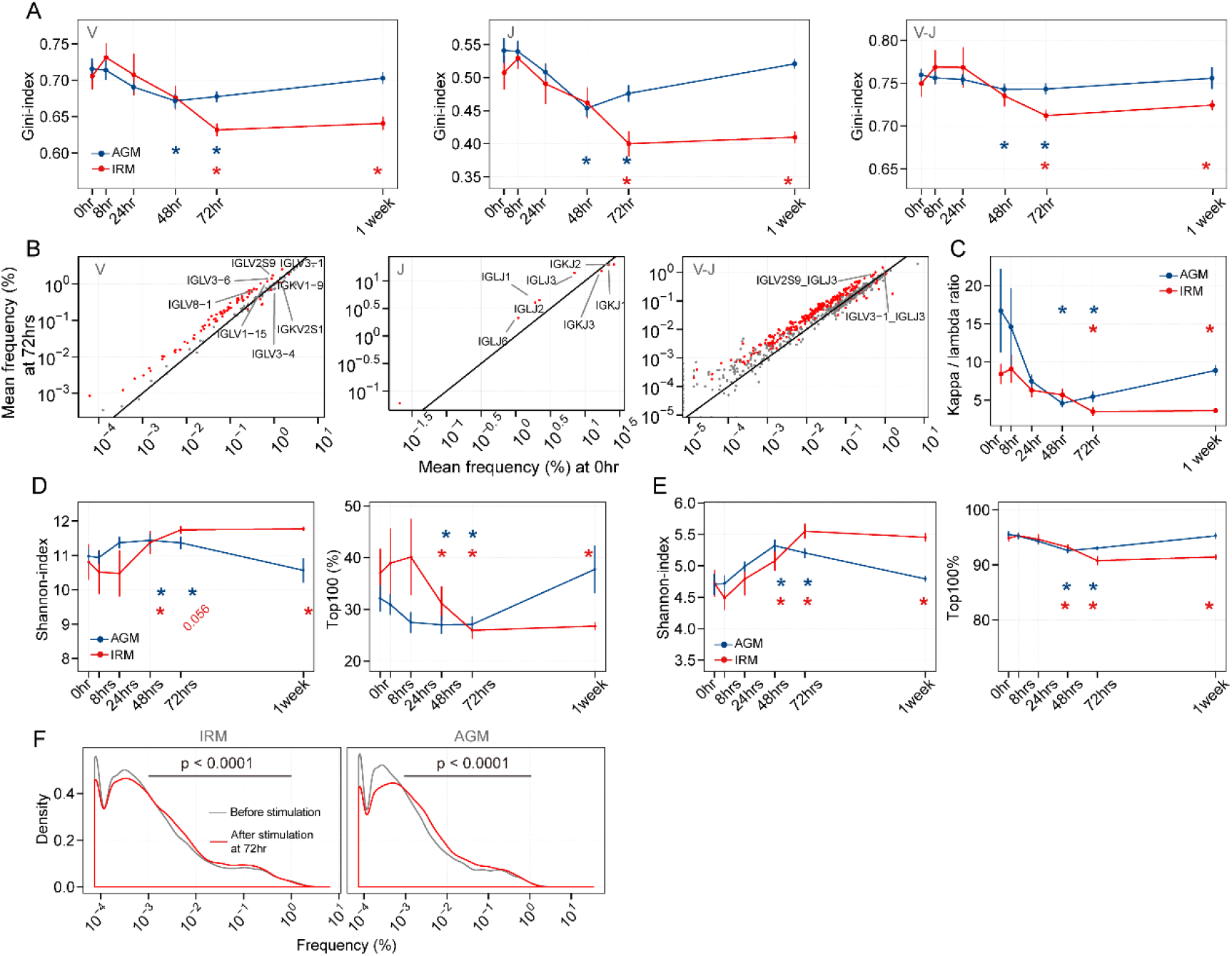
IgKL gene usage and repertoire diversity after the TLR7/8 stimulation in RMs and AGMs. (**A**) Gini-index of V-(left), J-(middle), and V-J genes pairings(right). (**B**) IgKL gene usage in RMs at 0hr and 72hrs. Mean frequency of the V-(left), J-(middle), and V-J pairings (right). Red dots are the genes with a p-value <0.05. (**C**) The κ to λ ratio of IgKL repertoire. (**D**) Shannon-index (left) and Top100 of clones (right). (**E**) Shannon-index (left) and Top100 of lineages. (**F**) The frequency distribution of lineages presented at both 0hr (grey) and 72hr (red) (Kolmogorov-Smirnov test was used). The paired Wilcox-ranked test was used in **A**-**E** (* p<0.05).

Then, we evaluated the diversity of BCR repertoire by Shannon-index and the percentage of top100 abundant clones. For AGMs, Shannon-index of IgKL clones gradually increased from 24hrs to 1-week in RMs, and then declined after 72hrs; For RMs, Shannon-index of IgKL clones sightly decreased from 0 to 24hrs, raised from 24hrs to 72hrs, and kept stable after 72hrs (**Fig. 2D**). RMs showed an obvious increase of Shannon-index at both 48hrs and 1 week, while AGMs presented obviously increased Shannon-index at both 48hrs and 72hrs. Meanwhile, reversed trends were presented for RMs and AGMs when the percentage of top100 was calculated (**Fig. 2D**). For top100, RMs exhibited obvious increases from 48hrs to 1 week, and AGMs showed obvious increases at both 48hrs and 78hrs.

To understand how TLR7/8 stimulation affected BCR repertoire function, we evaluated the characteristics of lineages. A lineage is a group of clones with identical V, J, and CDR3 amino acid sequences with no more than one mismatch. In theory, clones from a given lineage derive from an ancestry B cell^20^, and recognize the same antigen with different affinities. For AGMs, Shannon-index raised from 0 to 48hrs, and then decreased from 48hrs to 1 week; for RMs, Shannon-index of IgKL clones sightly decreased from 0 to 24hrs, raised from 24hrs to 72hrs, and keep stable after 72hrs. Reversed trends were presented for RMs and AGMs when the percentage of Top100 was calculated, suggesting that high frequent lineages reduced after the TLR7/8 stimulation (**Fig. 2E**). To identify the lineages affecting the BCR repertoire diversity, we compared the number of lineages with different frequencies at 0hr and 72hrs. We showed that, at 72hrs, the number of lineages with frequency <0.001% reduced, and the number of lineages with frequency 0.001% ~ 1% raised in both RMs and AGMs (**Fig. 2F**). Therefore, there is an expanded number of lineages with frequency 0.001% ~ 1% at 72hrs.

We also evaluated gene usage and diversity of IgH repertoire in RMs. Gini-index of V, and J genes usage raised from 0 to 24hrs, reduced from 24hrs to 72hrs, and kept stable after 72hrs. Shannon-index of clones and lineages decreased from 0 to 24hrs, reduced from 24hrs to 72hrs, and then became constant after 48hrs (**Fig. S2**). Top100 decreased after stimulation, peaked at 24hrs, declined from 24hrs to 72hrs, and then became stable. In conclusion, the TLR7/8 stimulation skewed IgH and IgKL repertoires in different kinetics.

In contrast, we stimulated these NHPs two months later with a TLR9 agonist, and examined the kinetics of BCR repertoire response under the TLR9 stimulation. Gini-index analyses showed a slight but gradual increase of the evenness of V, J, and V-J pairings usage in IgKL. (**Fig. S3A**). In RMs, Shannon-index of IgKL lineages reduced during one week; In AGMs, Shannon-index of IgKL lineages fluctuated from 0hr to 24hrs, and then significantly decreased from 48hrs to 1week (**Fig. S3B**). For IgH, Shannon-index reduced from 0hr to 48hrs, raised from 48hrs to 72hrs, and then kept stable until 1week (**Fig. S2D**). Therefore, we suggested that BCR repertoire of NHPs responded to the TLR7/8 stimulations and the TLR9 stimulation in different kinetics.

### The TLR7/8 stimulation expanded low-frequent but highly mutated lineages in NHPs

We further examined frequency, unique clones, and mutation rate (termed three features) of lineages. According to features respectively, we classified lineages into five classes, stable-lineage, increased-lineage, decreased-lineage, new-lineage, and disappeared-lineage (see **Methods**). Among them, increased- and new-lineages belonged to “Expanded-lineages”, and decreased- and disappeared-lineages belonged to “Shrinking-lineages”. In RMs, as for lineage frequency, unique CDR3 number, and mutation rate separately, Expanded-lineages was more than Shrinking-lineages from 8hrs to 72hrs. Then the difference between them kept stable (**Fig. S4A-C** and **Fig. S5A-C**). As for these three features in AGMs, Expanded-lineages were more than Shrinking-lineages from 8hrs to 48hrs. The difference kept stable from 48hrs to 72hrs, and declined at 1week (**Fig. S4 D-F** and **Fig. S5 D-F**). We further calculated the fold change (FC) of lineages’ features between an original at 0hr and a subsequent measurement at a given time (see **Methods**). As the FC increased for these three features in RMs, increased-lineages became more than decreased-lineages. This difference between increased-lineages and decreased-lineages raised from 8hrs to 72hrs and then kept stable (**Fig. 3A-C**). As the FC grew for these three features in AGMs, increased-lineages also became more than decreased-lineages. This difference between increased-lineages and decreased-lineages raised from 8hrs to 48hrs, kept consistently from 48hrs to 72hrs, and declined at 1week (**Fig. 3D-F**). For IgH of RMs, increased-lineages were slightly more than decreased-lineages when considering lineages frequency and mutation rate (**Fig. S6**). Additionally, as fold change of frequency and mutation rate increased, IgH increased-lineages became more than decreased-lineages. Their difference raised from 0hr to 72hrs, and then declined at 1week (**Fig. S7**). Notably, for lineages with frequency over 0.001% at 72hrs, increased-lineages, new-lineages and disappeared-lineages were more mutated than other lineages (**Fig. 3G**). Since the TLR7/8 stimulation could not induce somatic hypermutation without a combination of BCR signaling, the increased number, unique CDR3 number, and mutation rate indicated that the TLR7/8 stimulation expanded low-frequent but highly mutated lineages.

**Fig.3.**
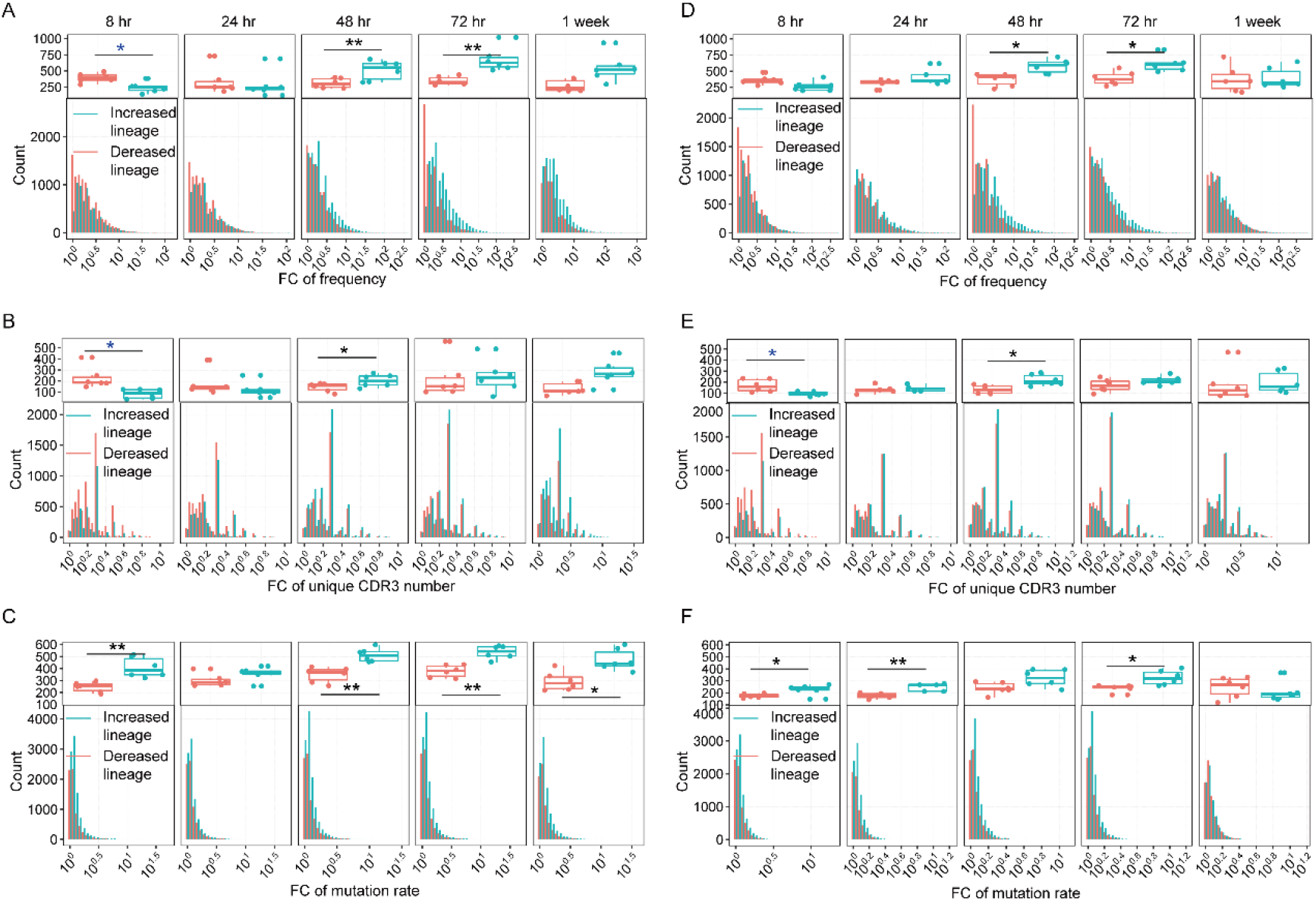
Fold change of three features of IgKL increased- and decreased-lineage distribution after the TLR7/8 stimulation. (**A**) Frequency FC distribution of increased-lineages and decreased-lineage in RMs. (**B**) Unique CDR3 number FC distribution of increased-lineages and decreased-lineage in RMs. (**C**) Mutation rate FC distribution of increased-lineages and decreased-lineage in RMs. (**D**) Frequency FC distribution of increased-lineages and decreased-lineage in RMs. (**E**) Unique CDR3 number FC distribution of increased-lineages and decreased-lineage in RMs. (**F**) Mutation rate FC distribution of increased-lineages and decreased-lineage in RMs. Upper plot of **A** and **D** showed the number of lineages with frequency FC>3. Upper plot of **B** and **E** showed the number of lineages with unique CDR3 number FC>2. Upper plot of **C** and **F** showed the number of lineages with mutation rate FC>1.3. The paired Wilcox-ranked test was used, and *p < 0.05, **p < 0.01.

### B cell activation was not necessary of lineages expansion

Since the TLR7/8 stimulation induced noticeable BCR repertoire changes at 72hrs, we set thresholds on three features at 72hrs to isolate expanded lineages (see **Methods**). In total, we identified 4,653 expanded lineages from six RMs, and 42.0% of these lineages met the thresholds on two or three features (**Fig. 4A**). As for the AGMs, we identified 3,941 expanded lineages, and 38.7% of them met the thresholds on two or three features (**Fig. 4B**).

**Fig.4.**
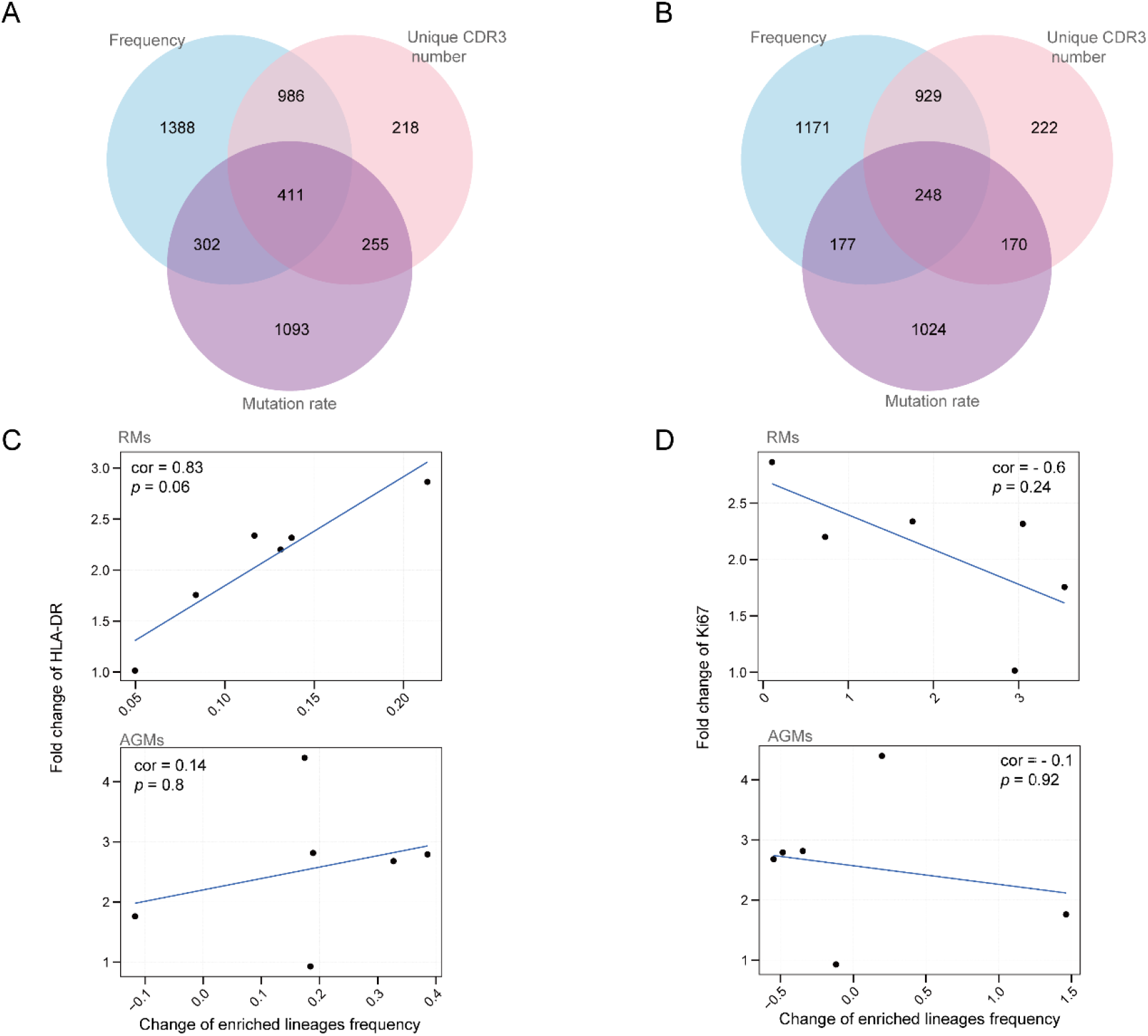
Identification of TLR7/8-expanded IgKL lineages and their association with B cell activation and proliferation. (**A**) Overlap of expanded lineages met thresholds on frequency, unique CDR3 number, and mutation rate in RMs. (**B**) Overlap of expanded lineages met thresholds on frequency, unique CDR3 number, and mutation rate in AGMs. (**C**) Spearman correlation between the change of expanded lineages’ frequency (%) and the change of B cell HLA-DR expression (%) in RMs (upper) and AGMs (lower). (**D**) Spearman correlation between the change of expanded lineages’ frequency (%) and the change of Ki67^+^ B cell (%) in RMs (upper) and AGMs (lower). The change of expanded lineages’ frequency (%) = (frequency at 72hrs - frequency at 0hrs)/ frequency at 0hrs; the change of HLA-DR MFI of B cells = (HLA-DR MFI at 24hrs - HLA-DR MFI at 0hr)/ HLA-DR MFI at 0hr.

A previous study shows that the TLR7/8 stimulation on purified B cells induces a lag secretion of antibodies after B cell activation^28^. We further examined the correlation between B cell activation and lineages’ expansion. For RMs, the fold change of sum-frequency of expanded lineages positively correlated (cor = 0.83) with the fold change of HLA-DR, and negatively correlated with the fold change of Ki67 (**Fig. 4C**). For AGMs, neither fold change of HLA-DR nor fold change of Ki67 correlated with the fold change of sum-frequency of expanded lineages (**Fig. 4D**). When considering unique clones only, the strong correlation between lineage expansion and HLA-DR expression was still robust in RMs (**Fig. S8**). These results suggested that B cell activation was not necessary of lineages’ expansion.

### TLR7/8-expanded lineages were more “private” than “public”

We further examined the characteristics of expanded lineages. The κ/λ ratio of expanded lineages was lower than random-sampled lineages in both RMs and AGMs. (**Fig. 5A**). 94.6% of expanded lineages using λ genes in RMs, and 93.5% in AGMs meet the frequency threshold (**Fig. s9**). It suggested that the TLR7/8 stimulation biased to expand lineages using λ genes, which raised κ/λ ratio of the entire BCR repertoire.

**Fig.5.**
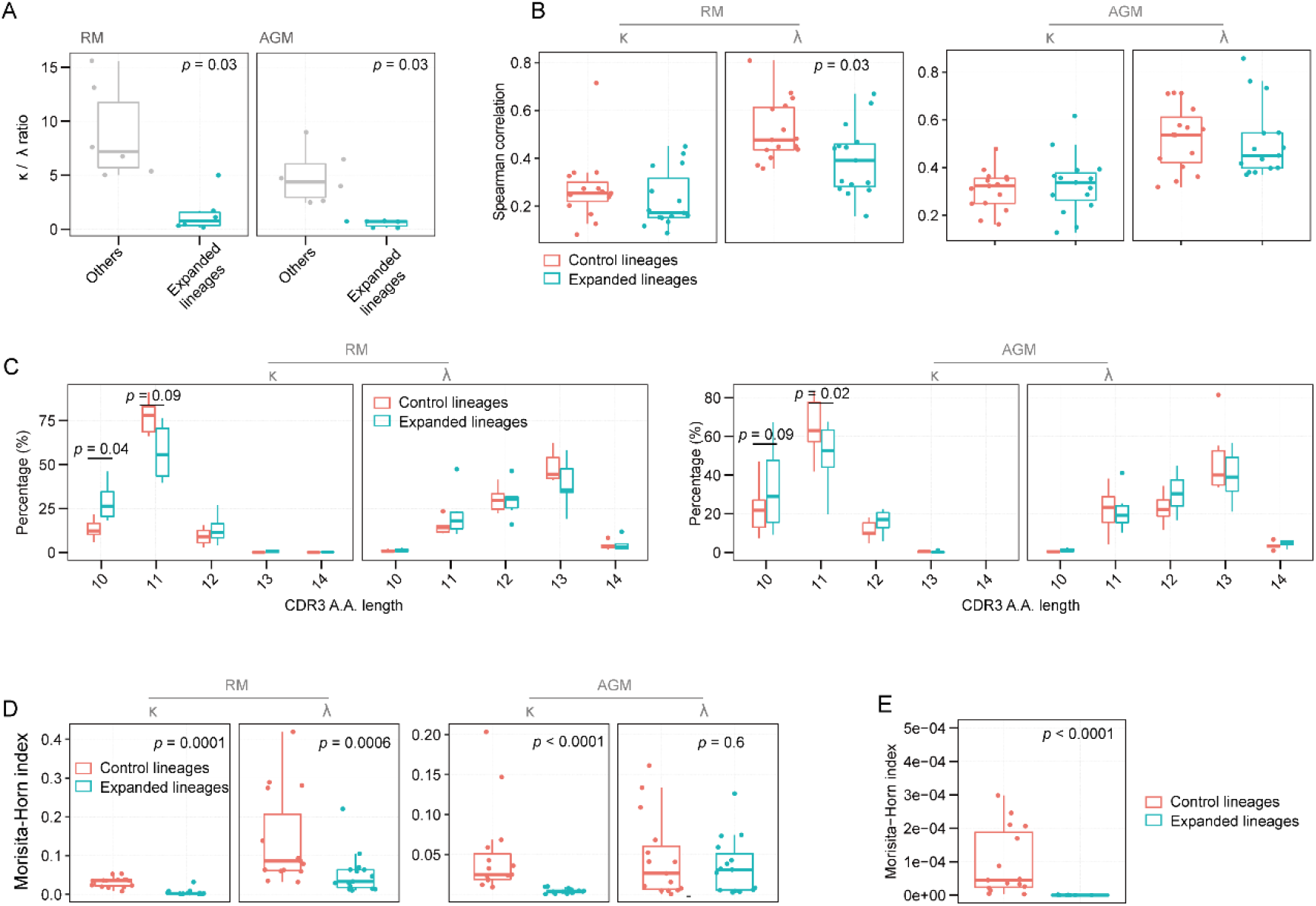
Characteristics of TLR7/8-enriched lineages at 72hrs. (**A**) The κ/λ ratio of expanded and Control lineages at 72hrs. (**B**) Spearman correlation of V gene usage of IgKL lineages between any two individuals. (**C**) The length distribution of CDR3 amino acid (A.A.) sequences from κ and λ IgKL lineages at 72hrs in RMs (left) and AGMs (right). (**D**) Intra-individual CDR3 similarity evaluated by Morisita-Horn index for IgKL expanded and Control lineages in RMs (left) and AGMs (right) at 72hrs. (**E**) Intra-individual CDR3 similarity evaluated by Morisita-Horn index for IgH expanded and Control lineages in RMs at 72hrs. Paired Wilcox-ranked test was used in (**A-E**).

**Figure 6.**
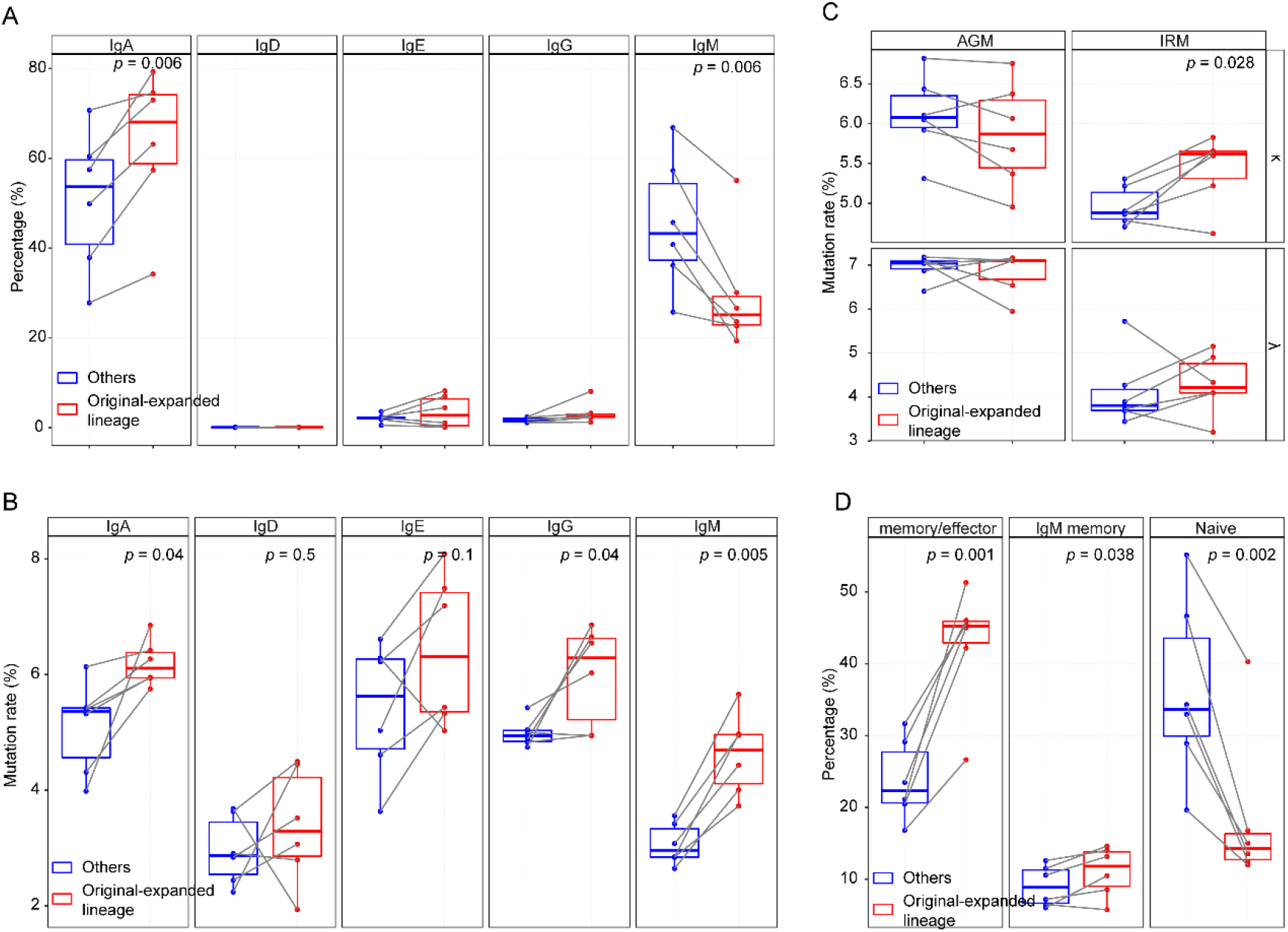
Mutation rate and isotype proportion of TLR7/8-expanded lineages at 0hr. We termed TLR7/8-expanded lineages at 0hr as Original-expanded-lineages, and other lineages at 0hr as Others. (**A**) Isotypes’ proportions of IgH Original-expanded-lineages and Others. (**B**) Mutation rate of each isotype of IgH Original-expanded-lineages and Others. (**C**) Mutation rate of IgKL Original-expanded-lineages and Others. (**D**) Proportion of memory, IgM memory and naïve B cells from which IgH Original-expanded-lineages and Others derived. Effector/Memory: IgA, IgG, and IgE clones with a mutation rate > 5. IgM memory: IgM clones with a mutation rate > 5. Naïve: IgM, and IgD clones with a mutation rate < 5. The paired Wilcox-ranked test was used for (**A-D**).

To examine whether the TLR7/8 stimulation would trig common lineages across subjects, we estimated gene usage, CDR3 length, and similarity of CDR3s of expanded lineages among subjects (termed interindividual similarity). We generated a control by randomly sampling IgKL lineages with the same κ/λ ratio and frequency distribution as expanded lineages (termed “Control” in following) (**see Methods**). For RMs, the interindividual similarity of κ V-/J-genes decreased, while interindividual similarity of λ V-/J-genes was similar in expanded and Control lineages; for AGMs, both interindividual similarities of κ and λ genes were equal between expanded and Control lineages (**Fig. 5B**). These results demonstrated that TLR7/8 stimulation didn’t drive a common gene usage, but induce a more heterogeneous gene usage among subjects. Secondly, we estimated the length distribution of CDR3s. As for clones using κ genes, CDR3s of expanded lineages in both RMs and AGMs were shorter than those of Control lineages; however, for clones using λ genes, expanded lineages had similar CDR3 length distribution as Control in both RMs and AGMs (**Fig. 5C**).

At last, we used Morisita-horn index to evaluate the interindividual similarity of CDR3 sequences^31^. For IgKL lineages, comparing with Control, similarities of either expanded κ lineages or expanded λ lineages reduced in RMs. The similarity of expanded κ lineages declined in AGMs (**Fig. 5D**). Additionally, interindividual similarity of CDR3 clones of expanded IgH lineages also declined (**Fig. 5E**). In conclusion, the TLR7/8 stimulation expanded “private” clones rather than “public” clones in peripheral blood.

### The TLR7/8 stimulation mainly expanded lineages from antigen-experienced cells

Class switching improves antibody affinity for antigens. We evaluated how the TLR7/8 stimulation affected isotype proportion. We randomly sampled an equal number of IgH lineages as a control. Expanded lineages consisted an increased percentage of IgA and decreased IgM comparing with control (**Fig. 7A**). At 0hr, IgA, IgG, and IgM clones of expanded lineages identified at 72hrs were higher mutated than those clones of control (**Fig. 7B**). For IgKL, expanded lineages were also higher mutated than control in RMs (**Fig. 7C**). According to the isotype and mutation rate of a lineage at 0hr, we inferred the state^29^ of cells expressing the expanded lineages, and classified them into three groups: memory/effector (lineages expressing IgA, IgG, and IgE), IgM memory^30^ (lineages expressing IgM with a high mutation rate) and naïve (lineages expressing IgD or IgM with a low mutation rate). Results showed that expanded lineages included a higher proportion of clones derived from memory/effector and IgM memory cells than control (**Fig. 7D**). Therefore, the TLR7/8 stimulation mainly activated BCR expression in antigen-experienced cells at 72hrs post-stimulation.

## Discussion

We showed that without a combination with antigen, the TLR7/8 stimulation alone could skew BCR repertoire in NHPs. This conclusion was supported by results that diversity and gene usage of IgKL and IgH repertoire were skewed after 8hrs post TLR7/8 stimulation, and an expansion of IgKL lineages happened in a time-dependent manner after stimulation. Some studies *in vitro* imply the activation of TLR7/8 may skew the entire BCR repertoire. John A Hanten *et al.* indicated that a ten-day TLR7/8 stimulation on purified B cells induced secretion of IgM and IgG of differentiated B cells^28^. Noa Simchoni *et al.* suggested that TLR7/8 agonist selectively expanded IgM^+^ and plasma cells expressing given IgH V and J genes^10^. This finding indicated that the TLR7/8 stimulation preferred to expand differentiated B cells, consistent with a higher TLRs expression level in memory B cells rather than in naïve B cells. Furthermore, it supported a view that TLR7/8 might bias to expand B cells expressing given BCRs. Our results develop this view, and identify that the TLR7/8 selection on BCRs can also happen *in vivo*. Other than memory IgM and IgG, IgA lineages in our study were also expanded by the TLR7/8 stimulation. Importantly, after the TLR7/8 stimulation, IgKL repertoire, which is overlooked in previous studies, showed a responding kinetic different to that of IgH repertoire. Similar to IgH repertoire results, high mutated IgKL lineages were the main subset expanded by the TLR7/8 stimulation. However, we found that the TLR7/8 stimulation biased to expand high mutated lineages with a low frequency. Furthermore, most of these expanded IgKL lineages were λ isotypes. Although we could not unveil the antigen-specificity of these expanded BCRs because of the technical limitations for identifying the antigens of BCRs, we showed that these expanded lineages had decreased interindividual similarity. This phenomenon suggested that the TLR7/8 stimulation in vivo expanded clones based on the status of cells expressing these lineages. Organ-derived clones were an origin of these expanded lineages. Genesis decides BCRs of residual B cells^31^. Differences in genesis among NHPs might induce different residual BCRs among individuals. In addition, NHPs experienced different antigen stimulations. B cells with a same status might express different BCR lineages among NHPs. At last, our results were slightly different from those presented by *in vitro* studies^10^. It was not a surprise because innate immune cells also expressed TLR7/8, and their participation might skew adaptive immune response.

TLR7/8-based adjuvants have a long history of efficacy in a pre-clinical and clinical setting. These studies highlight that TLR7/8 agonists promoted cellular immunity and antibody maturation. Previous investigations suggest that TLRs promote antibodies’ development by activating antigen-presenting cells (APCs) and B cells. However, high concentrations of numerous cytokines released by APCs and other immune cells can cause flu-like symptoms in humans. We showed that the TLR7/8 stimulation expanded low frequency but high mutated lineages. In contrast, the TLR9 stimulation on the same subjects expanded lineages with a higher frequency which induced a decreased IgKL repertoire diversity. Therefore, TLR7/8 signaling might enhance titers of antibodies just going through SHM when combining with an antigen stimulation. Importantly, similar to RMs, AGMs also expanded low frequency but high mutated lineages after the TLR7/8 stimulation, while lymphocytes activation were weaker in AGMs than in RMs. It suggested that the TLR7/8 stimulation was able to skew the BCR repertoire without over-activating immune cells. It will be valuable to study the intracellular signaling pathways of TLR regulating BCR expression and TLR regulating B cell activation in RMs and AGMs, because down-regulating TLR7/8 intracellular signaling pathways can reduce side effects of TLR7/8 adjuvants.

A notable finding is that the TLR7/8 stimulation biased to expand λ lineages in both RMs and AGMs. The decreased κ/λ ratio is typical of natural hosts of retroviruses. Our results indicate that ssRNA released by RNA viruses stimulates TLR7/8 in natural hosts, increasing λ chain usage in these primates. For heavy chain clones, class switching from IgD and IgM to IgG and IgA can increase clones’ affinity with the same CDR3 sequence for a given antigen. Lineages of λ and κ isotypes have different physicochemical and structural properties^32^. These differences may lead their different affinities for antigens^33^. Some studies have presented some evidence supporting this view. For example, antibodies from λ chains may dominate the neutralizing effect against SIV gp-120 in blood^34,35^. A recombination experiment showed that exchanging the light chain could reduce the binding affinity of antibody^36^. A study of the structure of an H3-clade neutralizing human monoclonal antibody suggests that light chains modulate the neutralizing spectrum of antibodies by affecting the local conformation of heavy chains^37^

A limitation in our study was the failure in sequencing IgH repertoire of AGMs. Ruijun Zhang *et al.^12^* identified some V and constant genes of AGMs by aligning genomics of AGMs^38^ to human reference adopted from IMGT. However, allelic diversity of genes and active SHM in IgH make it hard to find all references of AGMs IgH. In this study, we tried to design primers for constant, and V genes of AGMs based on reported references of both AGMs and RMs, but the alignment rate of AGMs IgH was less than 30% yet. The limited knowledge of AGMs BCR references also made it impossible to perform single-cell sequencing for AGMs BCR. AGMs have been identified as a more relevant model for studying human diseases and vaccination than mice^39–41^. Expanding the reference dataset of AGMs IgH will promote pre-clinical studies. To build this dataset, a large cohort of AGMs and long-read sequencing for the whole V-D-J regions enriched by 5’RACE method are needed in the future^42^.

## Acknowledgments

The authors would like to thank Kara S. Cox for performing flow cytometry.

## Author contributions

W.Z., I.M.W, X. L. designed and supervised the study; J.M., M.F. and M.C. collected samples; S.W. analyzed and interpreted the data; S.W. and W.Z. drafted the manuscript; I.M.W, X.L. L.L., Z.W., L.L., X.L., J.W., H.Y., N.S, T.L., C.N. and X.Z. revised the manuscript.

