## Supplemental figures for "Responding kinetic of B-cell receptor repertoire to the Toll-like receptor 7/8 stimulation in non-human primates"

**Supplementary Table 1.** Information of NHPs

| Group | Animal | Weight (kg) | Gender | Vaccination 1<br>18-Mar-2013<br>Time = 0 | Vaccination 2<br>13-May-2013<br>Time=8 weeks |
|---|---|---|---|---|---|
| IRM | A8R080 | 4.80 | F | IDR-053 (25mg) | IMO-2125 (2.5mg) |
| | A8R113 | 5.40 | F | | |
| | A8L111 | 6.75 | M | | |
| | A8R083 | 6.40 | F | | |
| | A8R066 | 5.80 | F | | |
| | A7E047 | 4.85 | F | | |
| | **avg wt** | **5.67** | | | |
| AGM | A8M004 | 5.65 | M | IDR-053 (25mg) | IMO-2125 (2.5mg) |
| | A8M017 | 5.15 | F | | |
| | A8M033 | 5.15 | F | | |
| | A8M042 | 4.70 | F | | |
| | A7M063 | 4.50 | F | | |
| | A8M010 | 6.30 | M | | |
| | **avg wt** | **5.24** | | | |

**Supplementary Table 2.** Constant region primers used in 5'RACE

| Primers | Isotype | Group |
|---|---|---|
| 5'-Bio ACACTTAATTAACCGGAGGTGGCATTGGAGG-3' | IgE | AGM |
| 5'-Bio ACACTTAATTAAC ATGGCGGGAAGATGAAGACA-3' | κ | AGM & IRM |
| 5'-Bio ACACTTAATTAAC AACGGAGTGACCGAGGGA-3' | λ | AGM |
| 5'-Bio ACACTTAATTAAC AACAGAGTGACTGACGGG-3' | λ | IRM |
| 5'-Bio ACACTTAATTAAC AACAGAGTGACCAAGGGG-3' | λ | IRM |
| 5'-Bio ACACTTAATTAAC AAVAGAGTGACCGWGGGG-3' | λ | IRM |

**Supplementary Table 3.** FACS antibodies

| Antigen | Clone | Fluorophore | Supplier |
|---|---|---|---|
| CD3 | 10D12 | Biotin/SA PercPcy5.5 | Miltenyi primary/BD secondary |
| CD4 | L200 | PE-CF594 | BD |
| CD8 | SK1 | PE-Cy7 | BD |
| CD20 | LT20 | VioBlue | Miltenyi |
| HLA-DR | L243 | APC-H7 | BD |
| CD69 | FN50 | FITC | BD |
| ki67 | B56 | PE | BD |

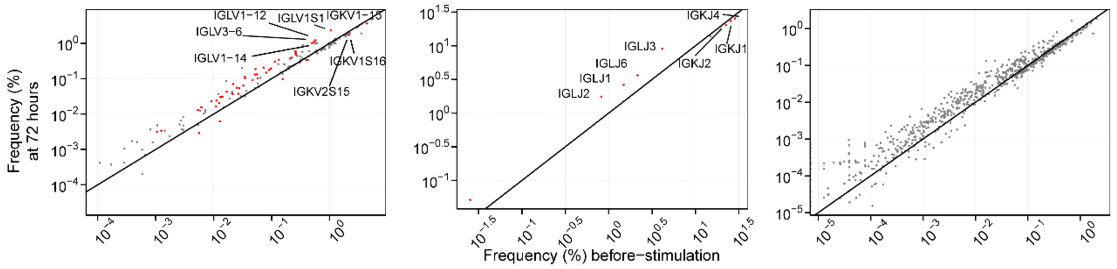


**Fig.S1 V-, J- and V-J paring usage at 0hr and 72hrs post-stimulation in AGMs.** The usage of V (left), J (middle) and paired V-J in AGMs before stimulation and after stimulation at 72 hours. Paired Wilcox-ranked test was employed to examine the difference of the gene usage, and then the p-values were corrected by FDR. The gene with corrected p-value < 0.2 were red.

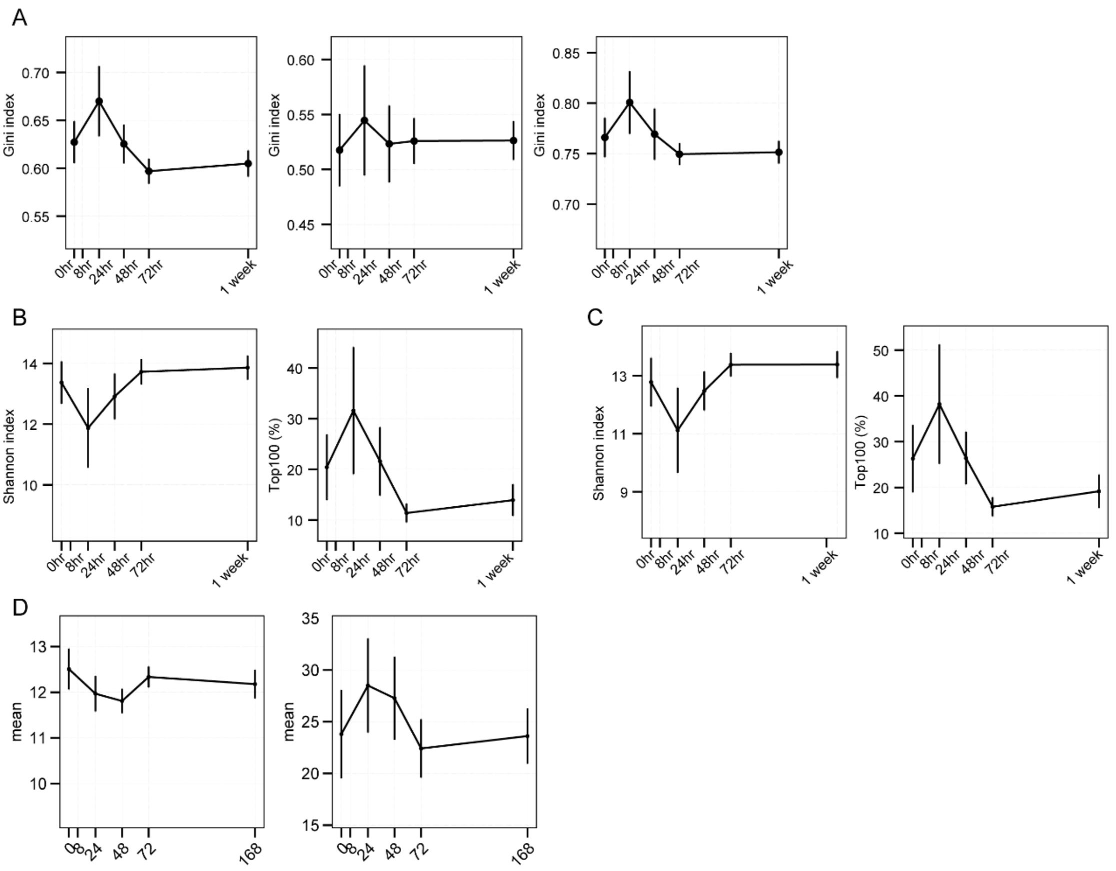


**Fig.S2 Responses of IgH to the TLR7/8 stimulation via TLR7/8 and TLR9 separately in RMs.** (**A**) Gini-index of V- (left), J- (middle) and V-J paring (right) in IRMs at 0hr, 24hrs, 48hrs, 72hrs and 1week post-stimulation via TLR7/8. (**B**) The Shannon index (left) and Top100 (right) of CDR3s for IRMs at 0hr, 24hrs, 48hrs, 72hrs and 1week post-stimulation. (**C**) The Shannon index (left) and Top100 (right) of lineages for each IRM before and after stimulation at 72 hours were presented in different colors. (**D**) The Shannon index (left) and Top100 (right) of lineages for IRMs at 0hr, 24hrs, 48hrs, 72hrs and 1week post-stimulation via TLR9. For (**A – D**), the paired Wilcox-ranked test was used, but no significant significance was observed.

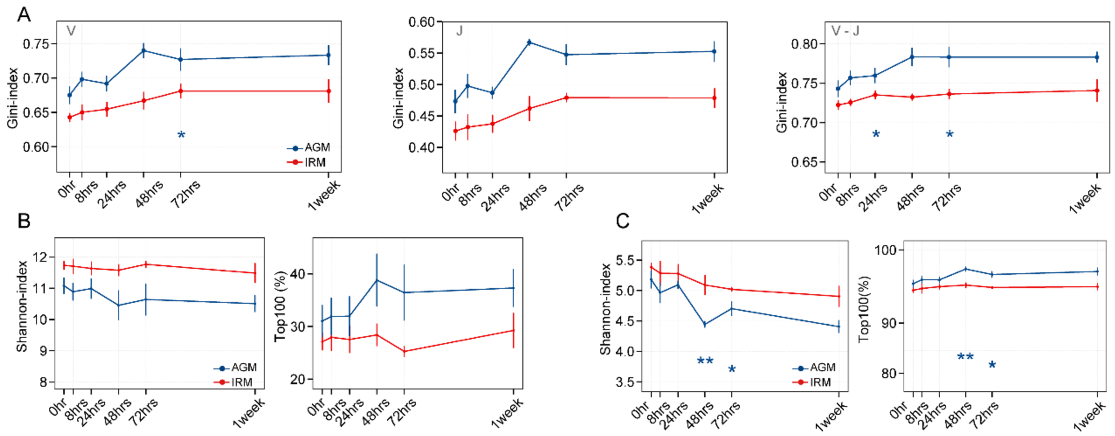


**Fig.S3 Analyses of the IgKL repertoire based on TLR9 agonist stimulation for both IRMs and AGMs.** (**A**) Gini-index of V- (left), J- (middle), and V-J pairings (right) of IgKL lineages. (**B**) Shannon-index (left) and Top100 of clones (right). (**C**) Shannon-index (left) and Top100 of lineages (right). Paired Wilcox-ranked test was used for a – e, *p < 0.05, **p < 0.01.

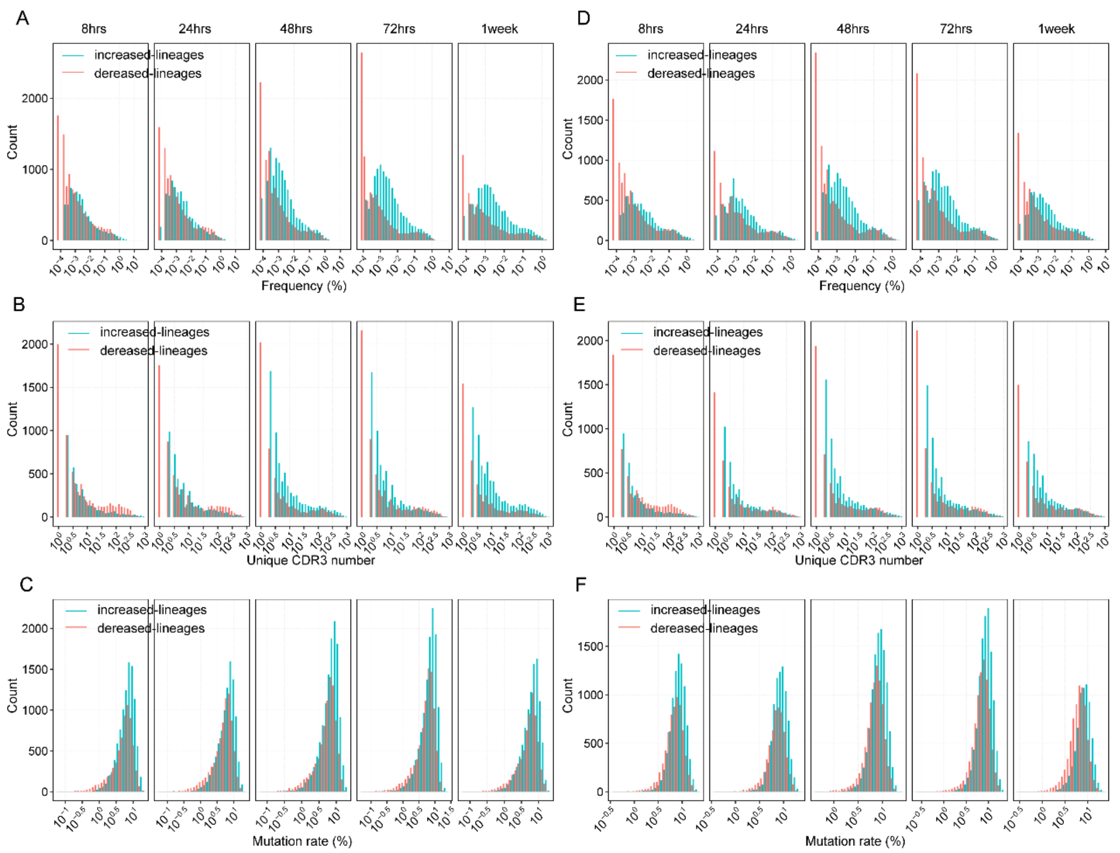


**Fig.S4 Distribution of frequency, unique CDR3 and mutation rate of IgKL increased-lineages and decreased-lineages from 8hrs to 1week in RMs and AGMs after the TLR7/8 stimulation.** The frequency distribution of increased-lineages (blue) and decreased-lineages (red) at 8hrs, 24hrs, 48hrs, 72hrs and 1week post-stimulation in RMs (**A**) and AGMs (**D**). The unique CDR3 distribution of increased-lineages (blue) and decreased-lineages (red) at 8hrs, 24hrs, 48hrs, 72hrs and 1week post-stimulation in RMs (**B**) and AGMs (**E**). The mutation rate of increased-lineages (blue) and decreased-lineages (red) at 8hrs, 24hrs, 48hrs, 72hrs and 1week post-stimulation in RMs (**C**) and AGMs (**F**).

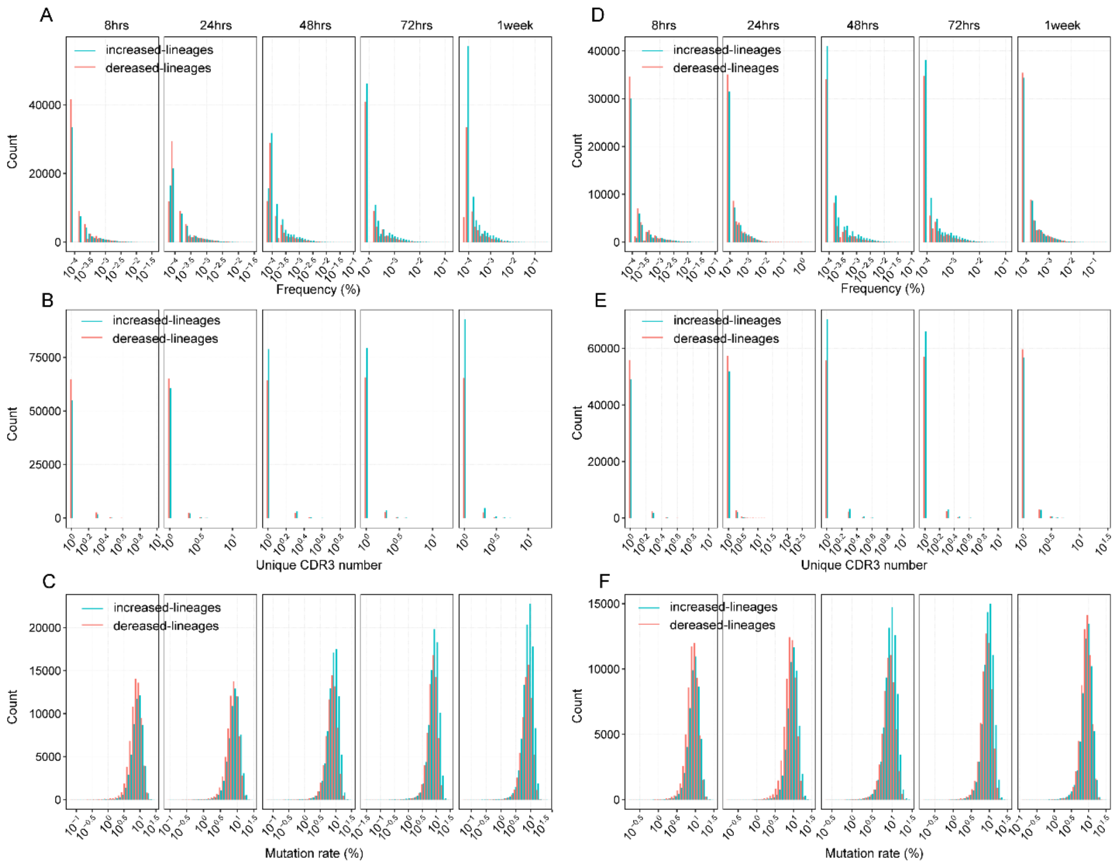


**Fig.S5 Distribution of frequency, unique CDR3 and mutation rate of IgKL new-lineages and disappeared-lineages in RMs.** The frequency distribution of new-lineages (blue) and disappeared-lineages (red) at 8hrs, 24hrs, 48hrs, 72hrs and 1week post-stimulation in RMs (**A**) and AGMs (**D**). The unique CDR3 distribution of new-lineages (blue) and disappeared-lineages (red) at 8hrs, 24hrs, 48hrs, 72hrs and 1week post-stimulation in RMs (**B**) and AGMs (**E**). The mutation rate of new-lineages (blue) and disappeared-lineages (red) at 8hrs, 24hrs, 48hrs, 72hrs and 1week post-stimulation in RMs (**C**) and AGMs (**F**).

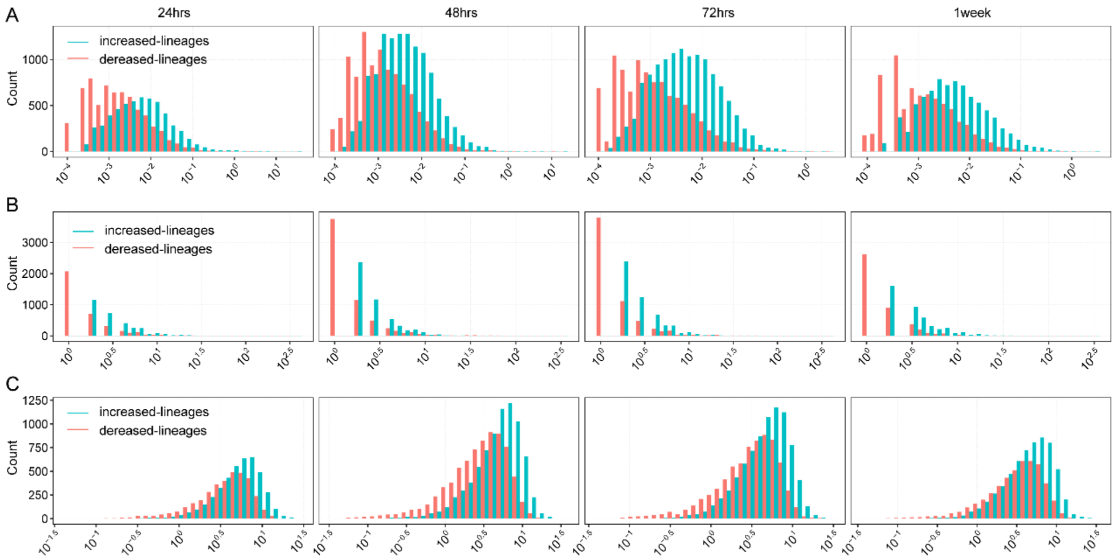


**Fig.S6 Distribution of frequency, unique CDR3 and mutation rate of increased and decreased lineages in heavy chain in IRMs.** The frequency distribution of increased-lineages (blue) and decreased-lineages (red) at 8hrs, 24hrs, 48hrs, 72hrs and 1week post-stimulation in RMs (**A**). The unique CDR3 distribution of increased-lineages (blue) and decreased-lineages (red) at 8hrs, 24hrs, 48hrs, 72hrs and 1week post-stimulation in RMs (**B**). The mutation rate of increased-lineages (blue) and decreased-lineages (red) at 8hrs, 24hrs, 48hrs, 72hrs and 1week post-stimulation in RMs (**C**).

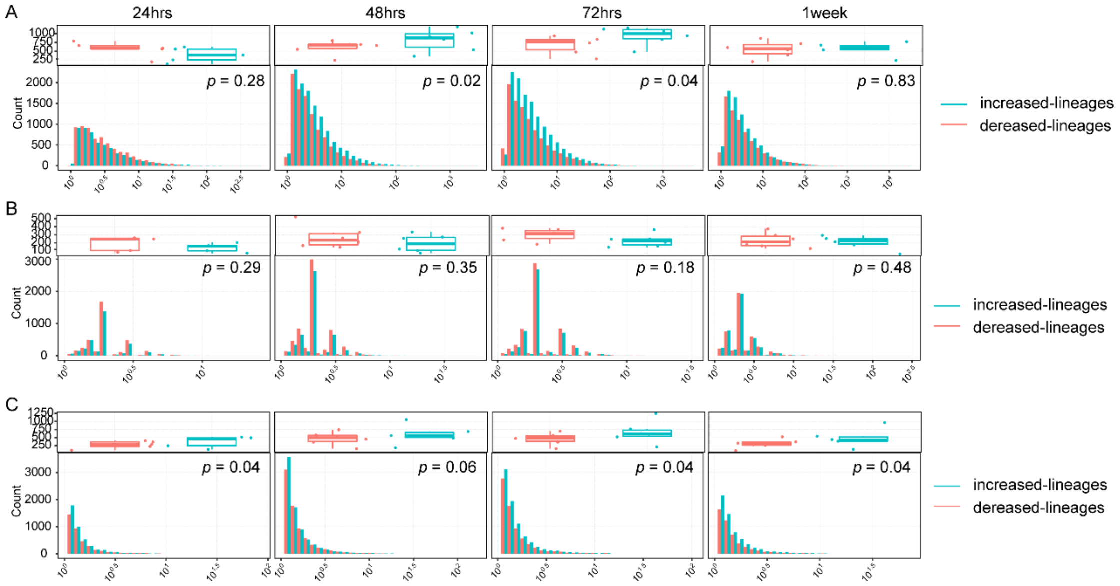


**Fig.S7 FC distribution of frequency, unique CDR3 and mutation rate of IgH increased-lineages and decreased-lineages in RMs after the TLR7/8 stimulation.** The frequency FC distribution of increased-lineages (blue) and decreased-lineages (red) at 8hrs, 24hrs, 48hrs, 72hrs and 1week post-stimulation in RMs (**A**). The unique CDR3 FC distribution of increased-lineages (blue) and decreased-lineages (red) at 8hrs, 24hrs, 48hrs, 72hrs and 1week post-stimulation in RMs (**B**). The mutation rate FC of increased-lineages (blue) and decreased-lineages (red) at 8hrs, 24hrs, 48hrs, 72hrs and 1week post-stimulation in RMs (**C**). For **A-C**, the paired Wilcox-ranked test was used.

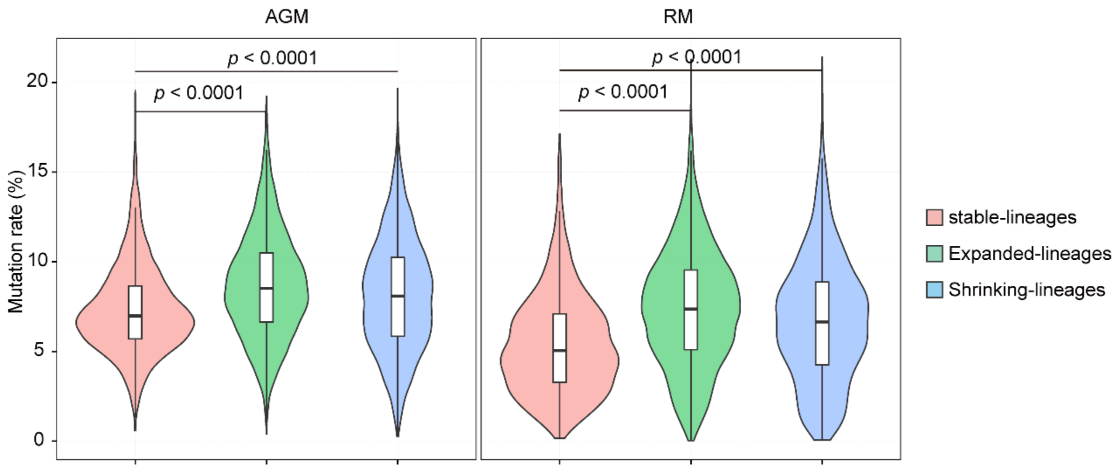

**Fig.S8 Mutation rate of lineages affected by the TLR7/8 stimulation at 72hrs.** The Wilcox-ranked test was used.

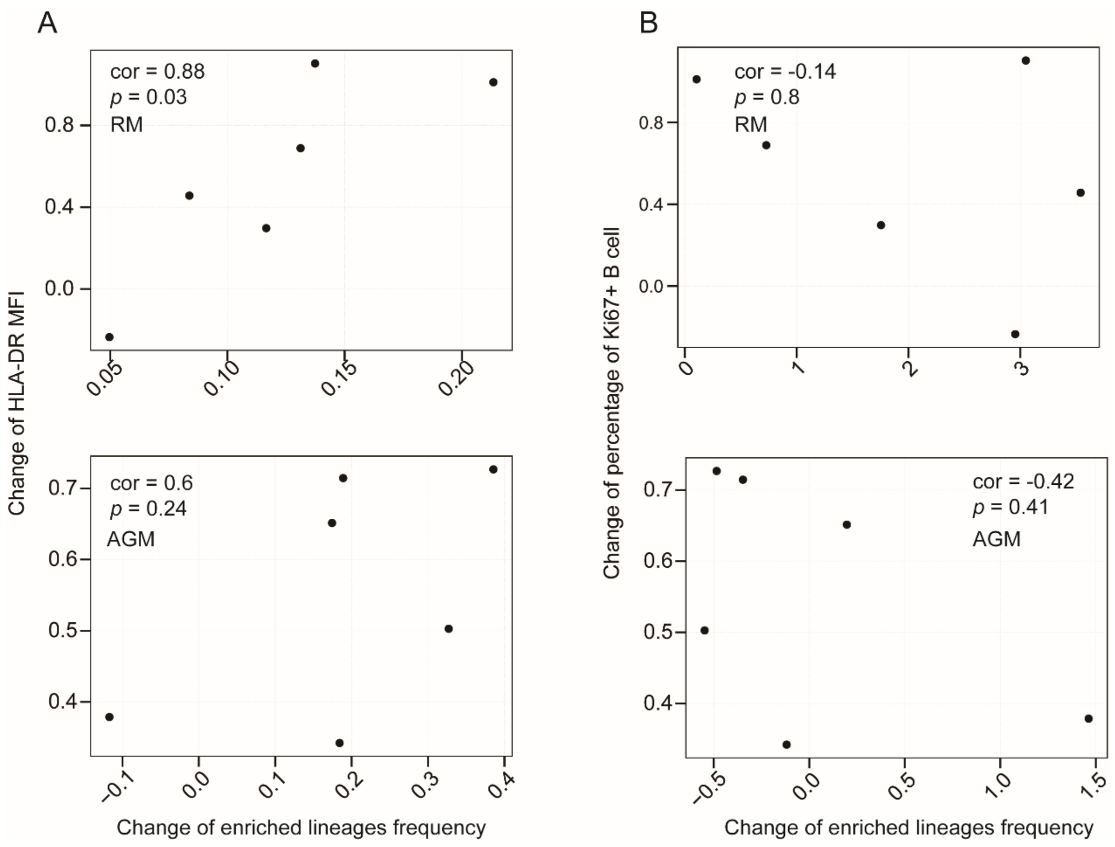


**Fig.S9 Correlation between increment of the number of enriched lineages and increment of HLA-DR expression and Ki67$^{+}$ percentage at 24hrs in RMs and AGMs after stimulation via TLR7/8.** Spearman correlation was employed.

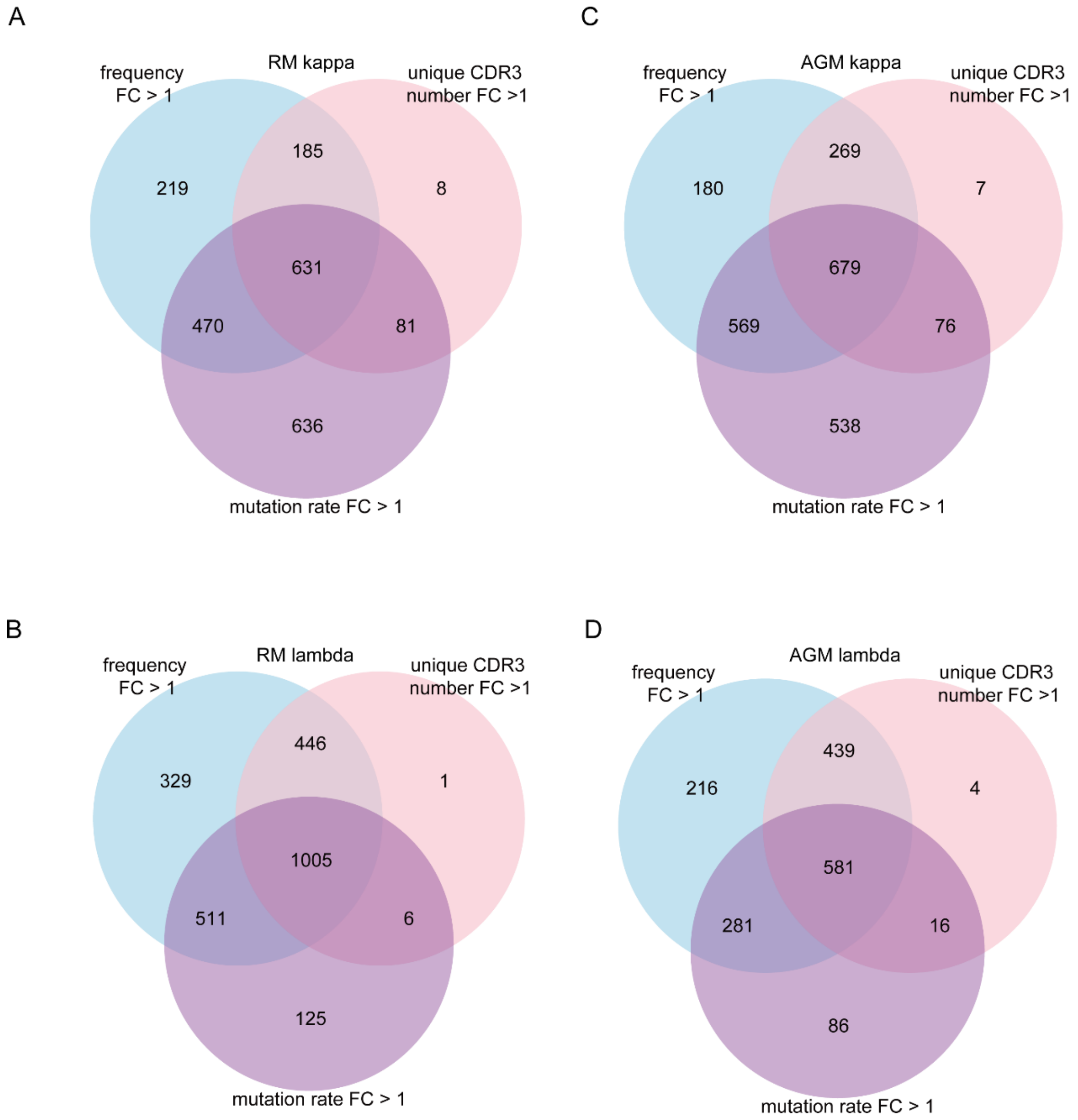


**Fig.S10 Number of expanded κ and λ lineages met thresholds on three features in RMs and AGMs. (A)** Expanded κ lineages met thresholds on three features in RMs. **(B)** Expanded λ lineages met thresholds on three features in RMs. **(C)** Expanded κ lineages met thresholds on three features in AGMs. **(D)** Expanded λ lineages met thresholds on three features in AGMs.
